# Unraveling the cis-regulatory code controlling abscisic acid-dependent gene expression in Arabidopsis using deep learning

**DOI:** 10.1101/2025.06.03.657584

**Authors:** Helder Opdebeeck, Dajo Smet, Sander Thierens, Max Minne, Herman De Beukelaer, Jasper Zuallaert, Michiel Van Bel, Marc Van Montagu, Sven Degroeve, Bert De Rybel, Klaas Vandepoele

**Affiliations:** Department of Plant Biotechnology and Bioinformatics, Ghent University, 9051 Ghent, Belgium; Center for Plant Systems Biology, VIB, 9051 Ghent, Belgium; Center for AI & Computational Biology, VIB, 9051 Ghent, Belgium; Department of Biomolecular Medicine, Ghent University, 9051 Ghent, Belgium; Center for Medical Biotechnology, VIB, 9051 Ghent, Belgium

**Keywords:** deep learning, transcriptional regulation, cis-regulatory elements, abscisic acid

## Abstract

Abscisic acid (ABA) is a key regulator of abiotic stress responses in plants. Understanding the regulation of ABA-dependent gene expression is key to uncovering how plants adapt to environmental stress and how their resilience can be improved. We explored gene expression regulation by ABA in *Arabidopsis thaliana* through the training of an interpretable deep learning model predicting ABA responsiveness in the root from proximal promoter sequences. Implementing state-of-the-art augmentation strategies to boost performance, our convolutional neural network-based model was able to confidently predict ABA responsiveness. We demonstrate that it learned actual motifs in the promoter sequences and confirm that ABRE-binding factor (ABF) binding sites play a key role in regulating ABA-dependent gene expression. However, also other motifs contributing to ABA-mediated gene expression regulation were identified. Our model outperforms a model trained to predict ABF binding, indicating it successfully learned the cis-regulatory code beyond the canonical ABF binding sites. Furthermore, the importance of motif clustering for regulating gene expression levels in response to ABA was unveiled by our model. Lastly, we identified genomic regions–beyond the proximal promoter–both with and without ABF binding sites, predicted by our model to drive ABA responsiveness. These genomic regions were used to generate reporter lines for experimental validation and were shown to drive the response to ABA *in planta*. This confirms that our model successfully inferred the regulatory code controlling ABA-dependent gene expression.

## Introduction

Understanding how plants regulate gene expression to adapt to their environment remains a major challenge in plant biology. Spatiotemporal control of gene expression is mainly governed by transcription factors (TFs), which recognize and bind short, specific DNA sequences known as TF binding sites (TFBSs) or motifs. Transcriptional regulation of gene expression is dictated by the availability of TFs and the number, position and organization of motifs, together comprising the cis-regulatory code (1). Understanding how plants coordinate transcriptomic responses to abiotic stress is becoming increasingly important in the face of ongoing climate change. Unraveling the mechanisms behind abscisic acid (ABA)-mediated gene regulation is crucial for understanding how plants cope with extreme abiotic perturbations, as ABA is a key regulator of abiotic stress responses and plays a pivotal role in triggering pre-existing adaptive strategies (2). The core ABA signaling pathway involves ABA binding to its receptors, leading to protein phosphatase inhibition and activation of protein kinases, which phosphorylate and activate ABRE-binding factors (ABFs). These bZIP TFs bind canonical ABF binding sites in non-coding DNA sequences to trigger the expression of neighbouring or distant ABA-responsive genes (3). However, many genes responding to environmental changes lack these known ABF TFBSs (4), indicating that other TFs are also involved and that our understanding of the cis-regulatory code, controlling how ABA regulates gene expression, is incomplete.

Computational approaches have long been central to studying transcriptional regulation in plants. Traditional methods such as co-expression (5) and gene regulatory network (GRN) analysis (6) have been applied to infer TF-target gene regulatory interactions. Feature-based machine learning (ML) methods have been used to unravel the cis-regulatory code underlying the regulatory interactions of a transcriptomic response. By training ML models on DNA motifs, as well as other DNA sequence features, the transcriptomic response to various stresses have been successfully predicted in several plant species (7–11). Explainable ML techniques allow model interpretation to identify the motifs or motif combinations on which a model relied the most to inform its predictions. While advantageous for its interpretability and ability to handle small datasets, feature-based ML relies heavily on feature engineering to achieve good performance. As an ML model learns only from the features extracted from a non-coding DNA sequence rather than from the full sequence itself, it may overlook subtle yet potentially causal elements of the cis-regulatory code embedded within the non-coding DNA.

Representation learning methods, in particular deep learning (DL), can overcome this limitation, as it has the ability to learn directly from non-coding DNA sequences. DL leverages hierarchical neural network architectures to automatically discover highly discriminative features, without the need for extensive feature engineering. Convolutional neural networks (CNNs) consist of multiple layers, each processing outputs from the previous layer to learn increasingly complex representations. CNNs are able to effectively uncover patterns and relations in non-coding DNA sequences, such as regulatory motifs and their combinations, that might be missed by feature-based ML. Early work by Washburn and coworkers (12) used CNNs to predict mRNA expression levels from whole-genome DNA sequence in maize and sorghum. More recently, Zhao and coworkers developed PlantDeepSEA, a deep neural network based tool to predict regulatory effects of genomic variants on chromatin accessibility and identify high-impact motifs within genomic DNA sequences of six plant species (13). Peleke and coworkers trained interpretable CNN models predicting gene expression profiles (low, medium and high expression) and extracting regulatory motifs from gene flanking DNA sequences for root and leaf tissues of four different plant species (14). Li and coworkers developed PhytoExpr, a DL framework–based on CNNs or a transformer–that predicts mRNA expression levels as well as plant species from proximal regulatory DNA sequences from 17 diverse plant species (15). Wang and coworkers developed the DL framework DeepCBA–combining multiple DL architectures, including CNNs–to predict gene expression in maize based on the genomic DNA sequence and chromatin interactions (16). These studies demonstrate the ability of DL to automatically extract complex and biologically relevant motifs from genomic DNA sequences and predict gene expression levels.

The regulatory motifs learned using this approach, however, lack specificity, as the underlying models are mostly predicting baseline gene expression levels. To understand ABA-mediated gene expression regulation, the cis-regulatory code controlling ABA responsive gene expression needs to be unraveled. The number of genes available for training a DL model is, however, significantly lower in gene response prediction (i.e., limited to responsive genes) compared to the prediction of gene expression levels (which includes all expressed genes), posing a major challenge. Here, we present a CNN-based DL framework that allowed us to accurately predict ABA responsiveness in the Arabidopsis root using proximal promoter DNA sequences only. Building upon the current state of the art of data augmentation and interpretable DL for genomic sequences, we identified known and novel motifs, uncovered insights into their patterning, thereby further deciphering the cis-regulatory code underlying ABA responsiveness.

## Results

### Predicting transcriptional gene expression responses based on promoter DNA sequences

To investigate the cis-regulatory elements underlying the ABA response in Arabidopsis, we modeled the transcriptional response of genes to ABA treatment using DNA sequences and convolutional neural networks (CNNs). Results from RNA-Seq transcript profiling on 3h ABA-treated Arabidopsis roots (17) were used to obtain log2 fold change (log2FC) values reporting differentially expressed (DE) genes between control and treatment. These ABA log2FC values were used to define two classes of genes based on their response patterns. Whereas the DE genes with highest log2FC values were used as the upregulated (UP) class, the non-responsive (NON) class was defined as genes not showing significant DE and discarding genes lacking expression in all replicates (Fig. 1A). To equally balance the classes and maximize the contrast with the UP genes, NON genes were undersampled, only retaining those with log2FC values closest to zero.

**Fig. 1.**
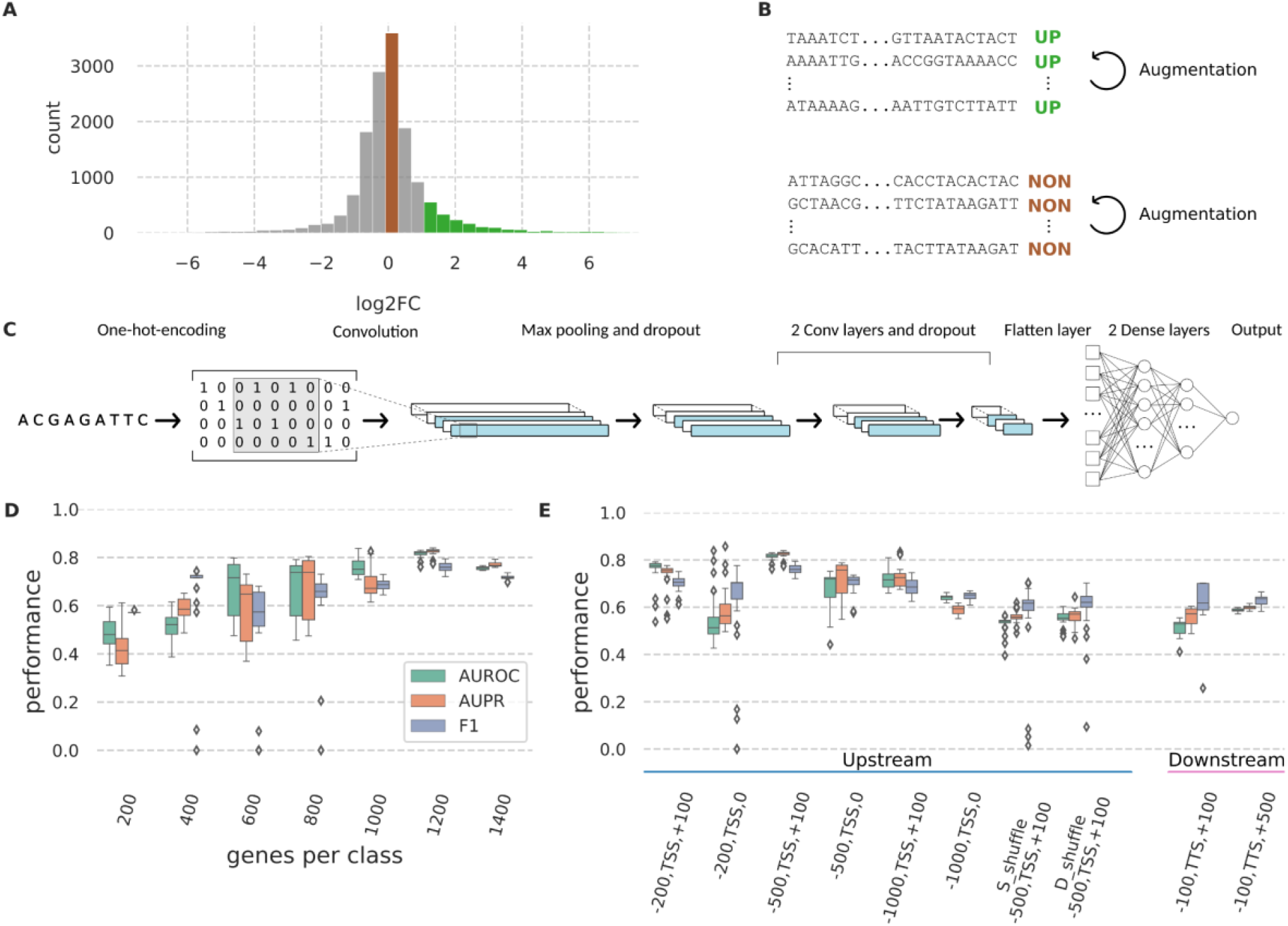
Data selection, training, and performance of the abscisic acid responsiveness deep learning model. (A) Distribution of Log 2 Fold Change (log2FC) values for differentially expressed genes after 3h of abscisic acid treatment in the Arabidopsis root. (B) Schematic overview of how the training set was assembled and expanded using different augmentation strategies. (C) Schematic overview of DL architecture for training. (D) Test AUROC (Area under the Receiving Operating Characteristic curve), AUPR (Area under Precision Recall curve) and F1 values for different balanced datasets. (E) Test AUROC, AUPR and F1 values for a range of different input regions. Blue indicates upstream input regions relative to the Transcription Start Site (TSS). Pink indicates downstream input regions relative to the Transcription Termination site (TTS) for the 1200 dataset. S_shuffle stands for single-nucleotide shuffle and D_shuffle for di-nucleotide shuffle.

Following balancing of the number of UP and NON genes, the data was divided into training, validation and test sets (80%, 10%, and 10%, respectively). In accordance with Washburn and coworkers (12), we made sure to avoid intra-gene family data leakage by partitioning our train, validation and test sets in such a way that genes from the same gene family were not shared between these sets. We used the area under the Precision-Recall and Receiving Operating Characteristic (AUPR and AUROC, respectively), as well as F1, calculated on the test set as performance metrics. Since the number of training samples (DNA sequences of UP and NON genes) is inherently limited, we employed several data augmentation techniques to enlarge the training set. First, we implemented reverse complement (RC) augmentation, in which each training instance was added once more with the same label, but as RC. Besides data augmentation *sensu stricto*, this form of augmentation enforced RC invariance at test time when using post-hoc conjoining (18). Secondly, we applied EvoAug, an augmentation framework that allows adding evolution-inspired augmentations (19) (Fig. 1B; *see Materials and Methods*). For the binary prediction problem, we used a CNN composed of three convolutional layers and two fully connected dense layers (Fig. 1C; *see Materials and Methods*). To account for the variability in model initialization and the subsequent effect on the test set performance, we always trained 20 different models on the same training set and reported median performance values.

The robustness of the models’ performance was assessed by varying the number of training genes and input regions. We initially used 600 base pair (bp) DNA sequences around the Transcription Start Site (TSS), defined as -500,TSS,+100 (i.e. 500 bp upstream and 100 bp downstream of the TSS) to optimize the number of training genes, ranging from 200 per class to 1400 (total number of ABA UP genes) per class. Genes were ranked according to their log2FC values. To reduce the number of training genes per class, we selected the UP genes with the highest log2FC values and an equal number of NON genes with the lowest absolute log2FC values. The ABA responsiveness models trained using 1200 genes per class yielded the best performance (median performance of 0.83 AUPR ± 0.02, a median AUROC of 0.82 ± 0.02, and a median F1 for the UP class of 0.76 ± 0.02). Despite the increase in contrast between classes, a smaller number of genes per class (stronger contrasts) did not lead to improvements in model performance but overall increased the variance, likely because models were trained on smaller datasets (Fig. 1D) (AUROC values of 0.48 ± 0.07 and 0.82 ± 0.02 for 200 and 1200 genes per class, respectively; *SI Appendix, Table S1*). Increasing the number of genes per class beyond 1200 reduced the classification performance. These results demonstrate that optimizing the number of training genes and the contrast between classes enhances the model’s performance in predicting ABA responsiveness.

Starting from these 1200 training genes per class, we assessed whether the chosen input region could be further optimized. We centered each input region either around the TSS for upstream sequences, or at the transcription termination site (TTS) for downstream sequences, and varied the size of these regions around these sites. Given the importance of the 5’ UnTranslated Region (UTR) for transcriptional regulation (9, 12, 14, 20), the model performance for upstream input regions with and without the first 100 bp of the 5’ UTR (median 5’ UTR length in the Arabidopsis genome is around 100 bp) was compared. The first 100 bp downstream of the TSS indeed proved important for classification performance, as excluding it, while keeping the region upstream of the TSS constant, led to a drop in performance (for example -500,TSS,100 compared to -500,TSS,0) (Fig. 1E). The optimal upstream region was determined to be 500 bp from the TSS (Fig. 1E). We also evaluated the classification performance on input regions from the terminator region. Both terminator definitions (−100,TTS,+100 and -100,TTS,+500) yielded performances around or slightly above random guessing (AUROC of 0.53 ± 0.03 and 0.59 ± 0.01, respectively). As a negative control, we randomized the DNA sequences of the best performing -500,TSS,+100 input region using mono- or dinucleotide shuffling. The observed drop in classification performance (AUROC 0.82 ± 0.02 to 0.54 ± 0.03 and 0.55 ± 0.03, respectively) for models trained on these shuffled sequences indicated that mono- or dinucleotide frequencies alone are insufficient for adequate classification (Fig. 1D, *SI Appendix, Table S1*).

We also compared the performance of our best-performing ABA responsiveness model with a classical feature-based machine learning approach, where we applied a random forest algorithm on a feature matrix reporting the presence or absence of known TFBSs in the -500,TSS,+100 sequences of UP and NON genes (11) (*SI Appendix, Fig. S1*). The DL model outperformed the feature-based ML approach, achieving an overall higher performance (AUROC 0.82 ± 0.02 for DL compared to 0.69 ± 0.0 for ML).

Finally, we retrained our CNN on a second dataset which quantified ABA responsiveness in Arabidopsis seedlings after 6h of ABA treatment (21). Using the top 1200 UP genes and a - 500,TSS,+100 bp input region, we also achieved a high performance (AUROC: 0.74, AUPR: 0.76, F1: 0.73), demonstrating our approach generalizes across diverse ABA responsiveness datasets.

### Different data augmentation strategies to improve the performance of DL models

To boost the success of training DL models, different augmentation strategies have been proposed. Apart from RC augmentation (18), the EvoAug framework enhances the training and generalization of genomic deep neural networks by applying a combination of evolution-inspired data augmentations (e.g. translocations, deletions, insertions, mutations, gaussian noise) (19). The more recent PhyloAug approach exploits the evolutionary sequence variation and conservation and proposes a phylogenetic augmentation approach by incorporating evolutionarily related genomic sequences from other species in the training set (22). Solely applying RC augmentation, we obtained a performance of AUROC = 0.69 ± 0.02 (AUPR = 0.70 ± 0.03, F1 = 0.67 ± 0.03). This strongly improved when applying EvoAug (AUROC of 0.82 ± 0.02, AUPR = 0.83 ± 0.02, and F1 = 0.76 ± 0.01) (Fig. S2A).

In an effort to better understand the effect of the different augmentation strategies offered by EvoAug, we applied each of the augmentations separately (*SI Appendix, Fig. S2B*). The mutation (AUROC = 0.71 ± 0.02) and gaussian noise (AUROC = 0.72 ± 0.02) augmentations resulted in a minor performance increase, while the deletion (AUROC = 0.80 ± 0.02), insertion (AUROC = 0.78 ± 0.03) and translocation (AUROC = 0.81 ± 0.01) strategies separately all resulted in a stronger performance increase. Nevertheless, the performance improvements for these separate augmentations were lower compared to the ensemble approach of EvoAug (median AUPR and F1 metrics are available in *SI Appendix, Table S2*). A shared characteristic of these augmentations is that they induce a shift in the sequence, which suggests this is one of the driving factors behind these successful augmentations.

To evaluate PhyloAug, we compiled a set of evolutionary related sequences from related accessions and species. Based on the PLAZA comparative genomics platform (23) and homology-based TSS annotation, 103,949 orthologous genes were derived from 7 other Arabidopsis accessions and 29,720 orthologous genes were derived from 8 other Brassicaceae species. These sets of genes were used in three different configurations: augmentation with promoter DNA sequences of orthologous Arabidopsis accession genes, orthologous Brassicaceae genes, and augmentation with the union of both (see *Materials and Methods* and *SI Appendix, Extended Methods*). Two parameters that are inherent to phylogenetic augmentation of a genomic dataset are the phylogenetic distance of the sequences and the augmentation rate. We determined the optimal combination of those parameters by performing a grid search (*SI Appendix, Fig. S2C*). Like this, we set the phylogenetic distance to maximum 4, which corresponds to creating the possibility of swapping out the training instances by orthologous sequences originating from either *Arabidopsis lyrata, Capsella rubbella* or *Cardamine hirsuta*. The augmentation rate was set at 0.5, which means that half of the training genes got swapped by orthologous sequences of one of those 3 species. Augmenting the training set with just the accessions had a minimal effect on the performance (AUROC = 0.69 ± 0.03, AUPR = 0.70 ± 0.03, F1 = 0.69 ± 0.02), while using the Brassicaceae species yielded a small improvement with an AUROC score of 0.72 ± 0.02 (AUPR 0.73 ± 0.02, F1 = 0.70 ± 0.02). Overall, these findings suggest that, within our ABA context, EvoAug enhances model performance more effectively than PhyloAug.

### Interpretation of the DL model identifies important regulatory motifs involved in ABA-dependent gene regulation

To investigate the ABA DL models’ capability to identify ABA-related regulatory motifs, we applied interpretation techniques focusing on the best-performing model comprising the top 1200 gene set and the -500,TSS,+100 region. We first applied DeepExplainer (24) to the trained model, obtaining nucleotide-level importance scores for the top 1200 UP gene sequences. These scores indicate the contribution of each nucleotide to the model’s classification decision. Next, TF-MoDISco (25) was employed to identify significant subsequences, termed seqlets, in regions with high-importance scores. TF-MoDISco clusters similar seqlets to generate patterns, represented as Contribution Weight Matrices, which are analogous to Position Weight Matrices used to represent TFBSs (25). By comparing the TF-MoDISco patterns to known plant TFBSs (see *Material and Methods*), we annotated TF-MoDISco patterns (Fig. 2A).

**Fig. 2.**
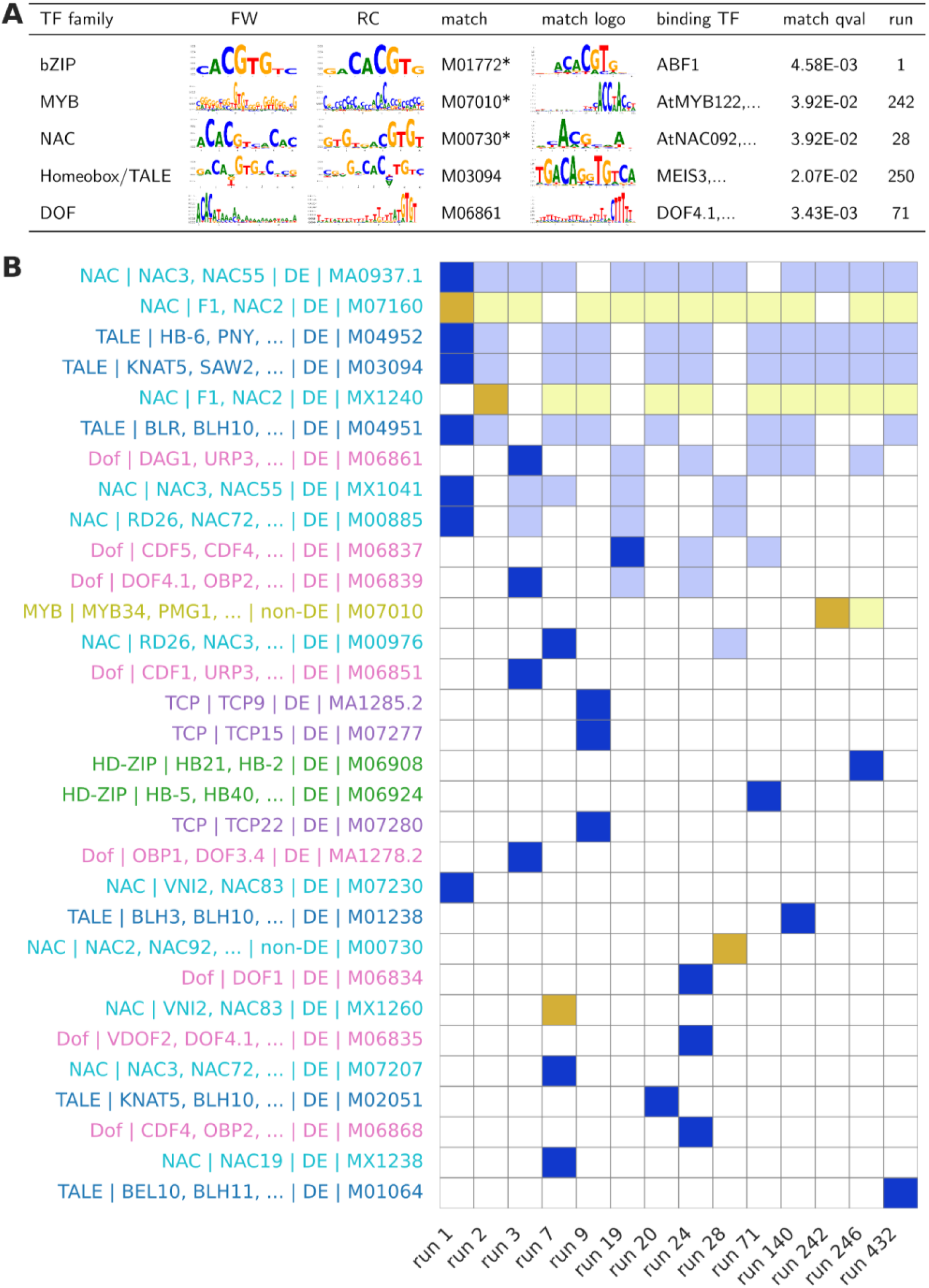
Deep learning model interpretation using TF-MoDISco. (A). Representative TF-MoDISco patterns across all ensemble runs. A representative significant known transcription factor (TF) binding site match is shown for each pattern. Asterisks denote Gold Standard (GS) motifs. (B) Illustration of the ensemble TF-MoDISco method. Only runs that exclusively discover patterns associated with TFs other than bZIP, bHLH or TriHelix families are shown here. For the full ensemble method output including runs that also recover bZIP, bHLH or TriHelix associated patterns, see SI Appendix, Fig. S3. Columns represent 14, out of 1278 total runs, sorted by the number of recovered known motifs. Rows show individual motifs and Y-axis label colors denote different TF families. Dark gold: GS known motif identified as TF-MoDISco pattern (q-value < 0.05). Light gold: GS known motif identified in an earlier run, with a higher number of recovered known motifs, and confirmed in the current run (q-value < 0.05). Dark blue: Non-GS motif with at least one binding DE TF identified (q-value < 0.05). Light blue: Non-GS motif identified in an earlier run, with a higher number of recovered known motifs, and confirmed in the current run (q-value < 0.05).

TF-MoDISco operates with several adjustable parameters to identify and cluster seqlets to delineate potentially important patterns. To ensure comprehensive pattern discovery and prevent important motifs from being overlooked due to limiting parameter combinations, we employed an exhaustive ensemble approach by running TF-MoDISco multiple times across a wide range of parameter values (*SI Appendix, Extended Methods)*. In total, we tested 1278 different parameter combinations (called “runs”, *Dataset S3*). We compared the set of significant TFBS matches over all runs to a set of 89 Gold Standard (GS) motifs, known to be involved in the regulation of ABA responses (*SI Appendix, Table S3*). GS recovery can be evaluated on two levels: the motif level and the TF family level. The performance of TF-MoDISco to identify GS motifs and TF families varied strongly depending on parameter settings (Fig. 2B, *SI Appendix, Fig. S3*). On the motif level, we counted how many GS motifs were found over all TF-MoDISco runs, from which we calculated the “motif recall” (i.e. recovery of ABA GS motifs). We next used the GS motifs to obtain a list of GS TF families and summarized the TF-MoDISco recall based on this list. To do so, GS motifs were grouped by the families of their associated TFs, forming “TF family motif groups” (n = 9). Then we checked how many of these GS TF family motif groups were associated with TF-MoDISco patterns. “TF family recall” is the proportion of “TF family motif groups” that are recovered (e.g. finding a NAC motif-like pattern with TF-MoDISco means the NAC TF family group was recovered).

Selecting the TF-MoDISco run that recovered most known TFBSs, we achieved a motif recall of 49%, a substantial improvement over the default settings which yielded a recall of 40% (representing a TF family recall of 33%, i.e. bHLH, bZIP, and NAC). By employing the ensemble approach we could further increase the recall: the motif recall reached 55%, while the TF family recall increased to 77%. Through this ensemble we identified motifs that corresponded to seven of the nine TF families part of the GS (ERF, HD-ZIP, MYB, NAC, ZF-HD, bHLH, and bZIP; Fig. S2). The motifs to which TFs of the CSD and CPP/TCR/CxC families belong, remained undetected. These TF families are, however, only represented by a single motif in the GS. Furthermore, our analysis identified additional patterns significantly matching known TFBSs that were not part of our GS. Forty-six of these motifs corresponded to DE TFs in our ABA dataset, including Trihelix TF GT-3a, BEL1-LIKE HOMEODOMAIN 1 and 8, and NAC55 and 83 (*SI Appendix, Fig. S3)*. These findings reveal that our interpretation analysis uncovered known motifs involved in ABA-mediated gene regulation beyond the GS. Moreover, comparing patterns matching known TFBSs from all ensemble runs (*SI Appendix, Fig. S4* and *Dataset S4)* revealed that while some TF families were easily found (e.g., bZIP, bHLH and BES family motifs were found in 55%, 68%, and 82% of all runs), motifs of other TF families were only found in a few runs (e.g., NAC and MYB family motifs were found in 11% and 0.4% of all runs).

### The importance of diverse TF motifs and cis-regulatory grammar to accurately predict ABA-dependent gene expression

A number of representative TF-MoDISco patterns (Fig. 2A), recovering a diversity of known TFBSs and TF families, were selected and highlighted on the visualized contribution scores for a selection of gene promoters (Fig. 3). The ABA responsiveness model classified the ABI FIVE BINDING PROTEIN 3 (*AFP3*, AT3G29575) proximal promoter as upregulated with the highest confidence (log2FC 4.0, prediction score 0.99). The bZIP pattern, identified 5 times by TF-MoDISco, matches significantly with the known ABF binding sites (Fig. 3). Similarly, in the proximal promoter of the *AWPM-19*-like gene (AT1G04560, log2FC 12.71, prediction score 0.98), encoding for a WPM membrane molecule, TF-MoDISco discovered two bZIP patterns significantly matching with known ABF binding sites (Fig. 3). WPM expression is known to be strongly induced by ABA treatment in wheat suspension-cultured cells, where the membrane protein helps to protect the cell against freezing (26). In rice, the AT1G04560 homolog PLASMA MEMBRANE PROTEIN1 (*OsPM1*), is directly regulated by the AREB/ABF transcription factor OsbZIP46, which binds to ABREs in the *OsPM1* promoter, activating its expression in response to drought stress (27). The proximal promoter of GALACTINOL SYNTHASE 1 (*GolS1*, AT2G47180, log2FC 2.0, prediction score 0.91), which encodes an enzyme involved in galactinol synthesis, contains an ABF as well as a MYB binding motif, which were recognized by TF-MoDISco (Fig. 3). GolS1 is a key osmoprotectant under various stress conditions (28) and is known to be regulated by Heat Shock Factors (29). In the promoter of *CYP94B3* (AT3G48520, log2FC 4.67, prediction score 0.86), a gene involved in the oxidation of the plant hormone jasmonoyl-L-isoleucine (30), TF-MoDISco identified patterns significantly matching DOF, NAC, Homeobox/TALE and bZIP binding sites (Fig. 3). *CYP94B3* contributes to the attenuation of the jasmonate signaling pathway (31) and is upregulated under ABA treatment, supporting the crosstalk between ABA and JA signaling (32). Finally, the patterns recognized by TF-MoDISco in the promoter of REVEILLE 6 (*RVE6*, AT5G52660, log2FC 1.72, prediction score 0.84), involved in the circadian rhythm, significantly match known DOF and bZIP bindings sites (Fig. 3). It has been shown that the ABA machinery is under circadian clock regulation which helps to modulate the stomatal opening during the day (33).

**Fig. 3.**
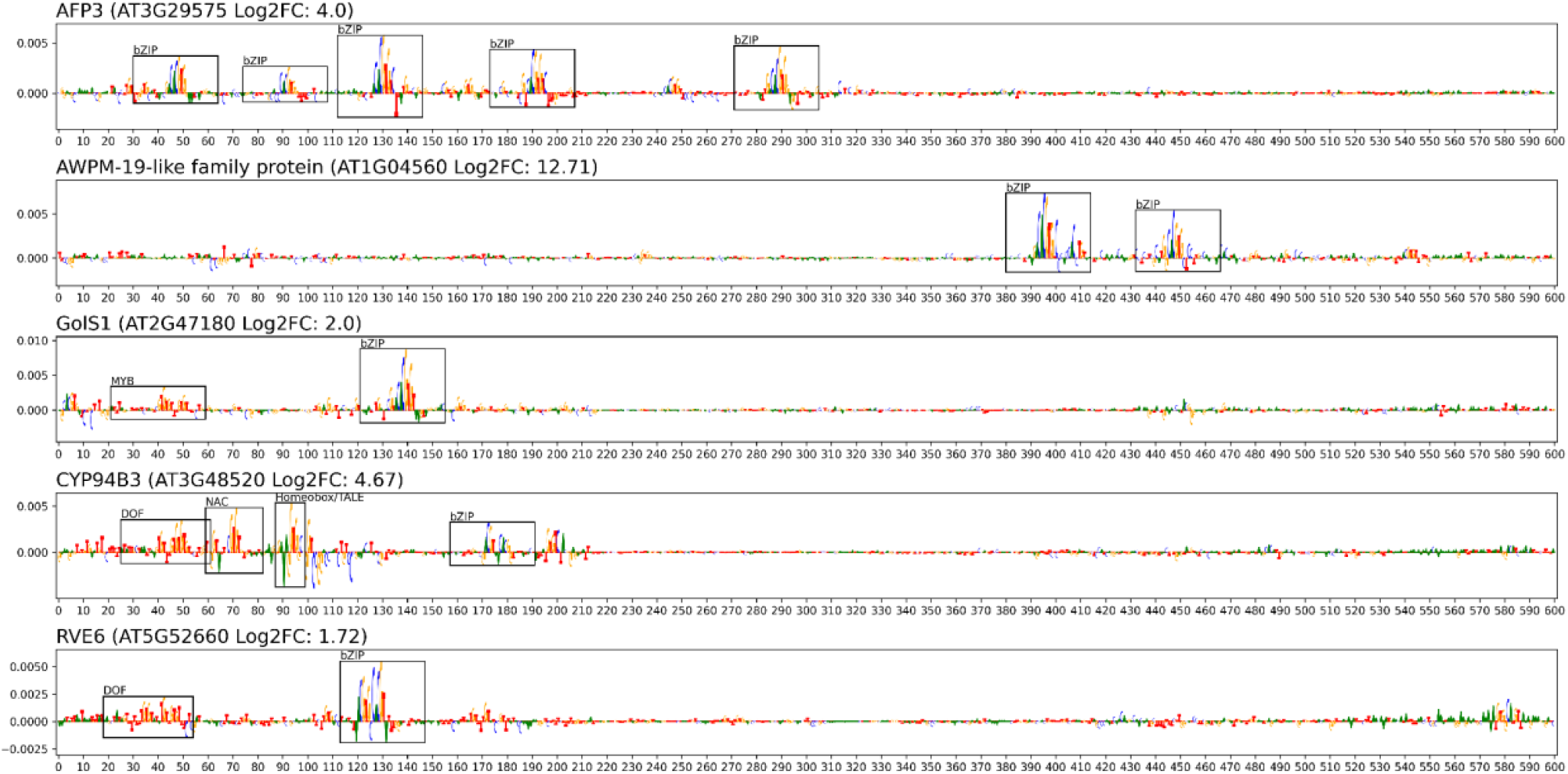
Cis-regulatory code learned by the deep learning model for a selection of abscisic acid upregulated genes. Selected DeepExplainer plots annotated with TF-MoDISco seqlets underpinning patterns from Fig. 2A. Plots show nucleotide-level importance scores (letter height) for predicting presence in upregulated class.

As these results corroborate the significance of ABF motifs while also suggesting that other motifs contribute to the model’s predictive performance, we next assessed how well our ABA responsiveness model performed compared to a DL model trained solely on ABF binding data.

Starting from 5726 high-quality ABF DNA affinity purification sequencing (DAP-Seq) peaks (17) and a similar number of non-peak regions, a DL model was trained to classify ABF peaks and non-peaks based on 600 bp DNA sequences (*SI Appendix, Extended Methods* and *Dataset S2)*. The resulting model could very accurately predict ABF binding (AUROC 0.98, AUPR 0.98, F1 0.96). Contribution scores obtained with DeepExplainer revealed a lot of G-box-like motifs, which explains the correct classification of regions containing ABF binding sites, and confirms previous results that members of the bZIP family, such as ABF, can bind to G-box-like motifs (34). Yet, 101 (6.7%) out of 1512 non-peak regions in the test set contained perfect G-box motifs (CACGTG) and 72 (71%) of these were nonetheless correctly classified as non-peaks. Furthermore, 316 (21%) and 344 (23%) of all 1512 non-peaks contained G-box-like motifs ACGTG and CACGT, respectively, and in both cases at least 85% of these were still correctly classified as non-peaks (*SI Appendix, Table S4*). This shows that the ABF binding model learned more than just the detection of G-box-like motifs.

Applying the ABF binding model to classify the promoters of the test set genes, used to evaluate the ABA responsiveness model, resulted in an AUROC of only 0.67 (AUPR 0.71, F1 0.53). This result indicates that the ABA responsiveness model (AUROC of 0.82, Fig. 1D) trained on promoter regions of UP genes was able to better learn the ABA cis-regulatory code than a model trained solely on ABF peak (and non-peak) sequences. This observation corroborates the large diversity in TF family patterns identified using the TF-MoDISco interpretation analysis (Fig. 2), indicating that on top of ABF motifs, also motifs from other TF families play an important role in controlling ABA-induced gene expression.

Besides recovering different motifs and TF families involved in ABA regulation, we next questioned whether the ABA responsiveness model also learned actual cis-regulatory grammar? Recent work showed there is a positive relationship between the number of binding sites for ABA responsive TF’s, a phenomenon called homotypic clustering (combinations of the same motifs), and the degree of ABA-induced expression (35). We first investigated the relationship between the number of seqlets corresponding to TF families detected by TF-MoDISco and the observed mean Log2FC of the 1200 UP genes. In agreement with previous findings, we found that having an increased number of bZIP motifs, here measured through ABF seqlets, was strongly associated with higher log2FC values (Fig. 4). Interestingly, this trend was also observed when evaluating the prediction scores for these genes, showing a maximum average prediction score > 0.75 for promoters with 3 ABF seqlets (Fig. 4). While this trend was also true for TALE motifs, the opposite pattern was found for B3 and C2H2 motifs. This highlights the importance of homotypic clustering of binding sites for certain TF families in modulating the strength of the expression response to ABA. Overall, while these results confirm that ABF TFs are key regulators controlling ABA-dependent transcription, the DL model identified additional motifs contributing to this transcriptional regulation.

**Fig. 4.**
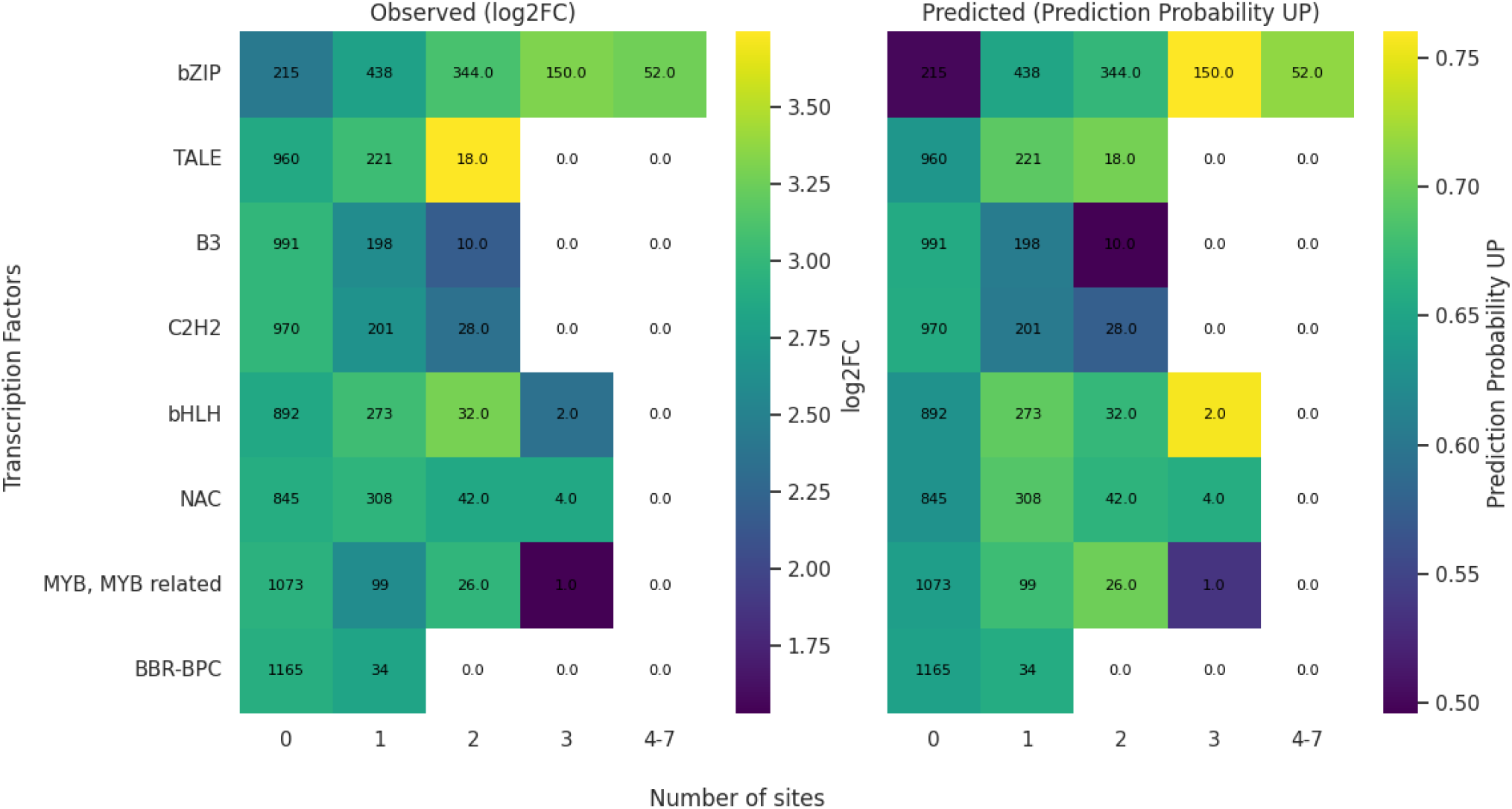
Effect of homotypic clustering of transcription factor family motifs on abscisic acid responsiveness and deep learning model prediction score. The x-axis denotes bins reporting the number of sites (seqlets of that transcription factor family) within a promoter. The y-axis denotes motifs from different transcription factor families. Values inside cells report the number of promoters. Color bars indicate the Log2 Fold change (log2FC) and prediction probability for being upregulated (UP).

### Experimental validation of cis-regulatory regions identified by the ABA DL model to drive ABA-dependent gene expression *in planta*

The global model performance and motifs identified using the interpretability analysis revealed that the ABA DL model successfully learned the cis-regulatory code controlling ABA-induced transcriptional responses. To assess the model’s ability to identify genomic regions involved in regulating the ABA response–beyond the proximal promoter–we used an ensemble of all 20 ABA responsiveness models as a robust predictor, averaging the models’ prediction scores. We initially focused on cases where the ABA UP regulation was incorrectly predicted. We identified 239 UP genes for which the prediction score for the 600 bp (−500,TSS,+100) proximal promoter was below 0.5, indicating a false negative prediction. For these genes, we next defined multiple 600 bp windows by sliding over the locus and computed the prediction score using the ensemble of DL models for each window (*SI Appendix, Extended Methods*). Focusing on the complete upstream sequence per locus, but excluding coding sequences, we selected the window with the highest prediction score and computed the score difference between this best-scoring window and the proximal promoter (called deltaS). We evaluated 53 loci with an upstream best-scoring window with deltaS ≥ 0.3 for ABF1–4 TF binding using DAP-Seq data (17) and found perfect agreement between these best-scoring windows and ABF peaks for 27 (51%) loci (*SI Appendix, Table S5*). This represents a 13-fold enrichment compared to randomly chosen control regions showing low DL prediction scores (prediction score best-scoring window < 0.2; 1/24 or 4% of these regions contain an ABF peak). Examples of ABA UP genes with distal best-scoring windows confirmed by ABF binding include DROUGHT-RESPONSIVE RING PROTEIN 1 (AT5G55970; best-scoring window 1.9kb upstream) and PHOSPHATIDYLINOSITOL-4-PHOSPHATE 5-KINASE 1 (*PIP5K1* or AT1G21980; best-scoring window 2.6kb upstream), as well as the NAC transcription factors *ATAF1* (AT1G01720) and *NAC083* (AT5G13180). While the former showed a very high deltaS value (0.61), the latter showed a distal best-scoring window 11kb upstream of the TSS.

As the ensemble of DL models identified for numerous misclassified ABA UP loci that the most important regulatory region is actually beyond the proximal promoter, we performed *in planta* experimental validation using reporter lines to assess whether these distal upstream regulatory regions can drive ABA-dependent transcription. Thirteen false negative ABA UP genes and their best-scoring distal promoter window were selected based on the prediction scores of the proximal promoter and the best-scoring distal window (using DeltaS, see above), the distance of the best-scoring window to the proximal promoter, and the presence of ABF peak(s) (*SI Appendix, Table S6*). Of these, eight had ABF peak support while five did not. Transgenic transcriptional reporter lines were generated by transformation, integrating the selected best-scoring window together with a minimal promoter and enhancer, and a β-glucuronidase (GUS) reporter in the Arabidopsis genome (see *Materials and Methods*). Reporter gene expression was investigated *in viv*o upon ABA treatment and compared to mock treatment (Fig. 5). Six out of eight best-scoring windows containing an ABF peak drove GUS expression in the primary root in response to ABA (Fig. 5 and *SI Appendix, Fig. S6*). The best-scoring window of *PIP5K1* (AT1G21980) induced expression in the columella, and the endodermis, cortex and epidermis of the elongation zone in response to ABA, with a strong induction in the stele. ABA induced the expression in the columella, and the stele of the elongation zone for the best-scoring window of AT4G30460. Expression was induced by the best-scoring window of PHYTOCHROME INTERACTING FACTOR 3 (*PIF3*; AT1G09530) in response to ABA in the columella, and strongly in the cells of the elongation zone (Fig. 5). ABA induced the expression in the quiescent center and other cells of the meristematic zone for the best-scoring window of RGA TARGET 1 (*RGAT1*; AT1G19530). Expression was induced by ABA in the columella, the cells of the meristematic zone and the stele of the elongation zone for the best-scoring window of AT4G19390 (mild expression in the stele also in the absence of ABA), whereas for that of *E12A11* (AT1G18100), ABA only induced expression upon lateral root initiation. Hence, the upstream regulatory regions - containing an ABF binding site - identified by the ensemble of DL models, confer ABA responsiveness in the root, primarily within the vasculature of the elongation zone.

**Fig. 5.**
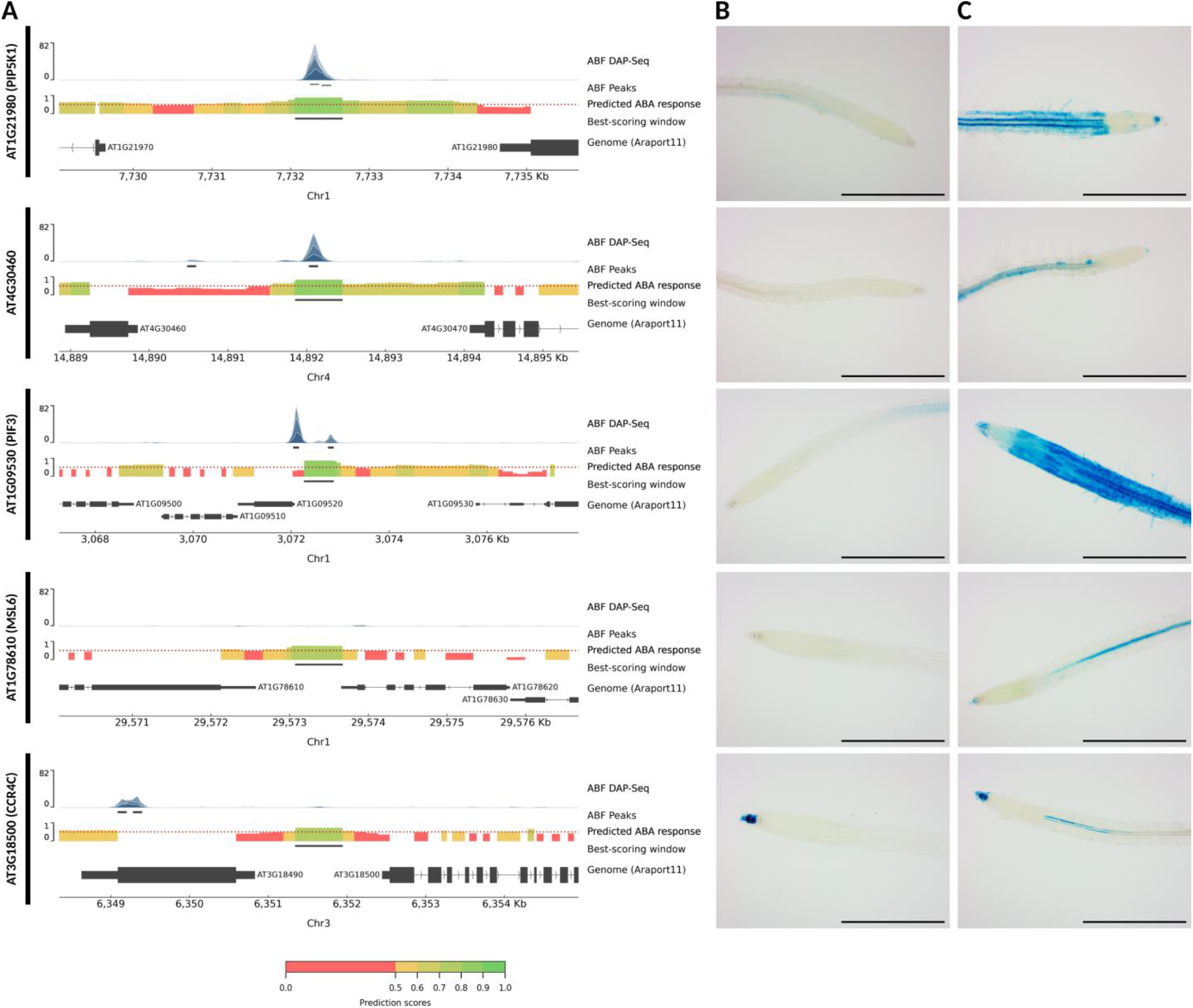
Experimental validation of ensemble deep learning model predictions using transcriptional reporter lines. (A). Overview of non-coding DNA regions–scanned with a window of 600 bp and stride of 300 bp–and corresponding ensemble model predictions. From top to bottom, the top track shows the presence of ABRE-binding factor (ABF) peaks (based on DAP-Seq data), with black lines indicating significant peaks. The middle track displays the median prediction score by the ensemble of DL models for each non-coding 600 bp window (color bar at the bottom of the figure), with a black line indicating the best-scoring window. A prediction score > 0.5 is indicative of an abscisic acid (ABA) responsive regulatory region. The bottom track shows the false negative gene of interest, neighbouring genes and their location in the genome. (B) and (C) In planta GUS reporter gene expression driven by the best-scoring window of a false negative gene of interest fused to a minimal promoter and enhancer in response to mock (B) and ABA treatment (C). The scale bar indicates a length of 0.5 mm. At least three independently transformed lines (n ≥ 3) were screened per treatment (mock and ABA) and per construct (e.g. best-scoring window).

Two of the five best-scoring windows lacking an ABF peak were also able to drive expression in the primary root in response to ABA. The best-scoring window of MECHANOSENSITIVE CHANNEL OF SMALL CONDUCTANCE-LIKE 6 (*MSL6*; AT1G78610) showed induction in response to ABA in the columella, and stele of the elongation zone, whereas that of CARBON CATABOLITE REPRESSOR PROTEIN 4 (*CCR4C*; AT3G18500) was only induced in the stele of the elongation zone (strong expression in the columella also in the absence of ABA) (Fig. 5). While screening ABA UP genes with low proximal promoter prediction scores, we also found examples where the best-scoring window was located in an intron. For *E1A1* (AT1G21400), our ensemble of DL models identified an intronic region for which the prediction score (0.807) was considerably higher compared to its proximal promoter (0.586). Similar to the distal upstream regulatory regions described above, we also generated a transgenic transcriptional reporter line, to assess whether this best-scoring intronic region can drive the ABA response. Reporter gene expression was induced by this intronic region in response to ABA. As the ensemble of DL models was trained on ABA UP genes in the root, we expected the expression to be induced at least in the root. Surprisingly, however, ABA-induced expression was observed in the leaf, not the root (Fig. S6), suggesting the presence of subtle tissue-specific regulatory elements that were not captured by the ensemble of DL models.

Taken together, these results suggest that our ensemble of DL models is able to identify cis-regulatory sequences beyond the proximal promoter that are biologically relevant and are able to drive an ABA-dependent transcriptional response. Furthermore, the ability of non-proximal promoter sequences without ABF binding sites–identified by our ensemble of DL models–to drive the ABA response, again indicates that it learned ABA-specific cis-regulatory elements beyond the canonical ABF-binding sites.

## Discussion

In this study, we presented a DL framework to predict and explain the ABA-dependent gene expression in the Arabidopsis root. We demonstrated the ability of our model to accurately predict ABA responsiveness, effectively learning the underlying cis-regulatory code. Besides canonical ABF binding sites, other TF families and binding sites involved in the regulation of ABA responsiveness were successfully identified. Applying our model as predictor we identified ABA regulatory regions in the Arabidopsis genome–beyond the proximal promoter used for model training–both with and without ABF binding sites, able to drive ABA responsiveness *in planta*.

In contrast to other plant DL frameworks (12–16), we used DL not to predict baseline gene expression but to predict ABA responsiveness. Unraveling the cis-regulatory code controlling ABA-dependent gene expression came with two major challenges: (1) To explain how ABA-dependent gene expression is regulated using interpretable DL, required a model accurately predicting ABA responsiveness (2) To train a model able to predict ABA responsiveness, only ABA-responsive (UP) genes, and an equal number of nonresponsive genes (NON) could be used. Because of these challenges, we could not build upon state-of-the-art deep neural network (DNN) based models, such as DeepSEA, DeepBind, Basset, Basenji, Borzoi and Malinois (36–38). These models typically have millions of parameters across hundreds of stacked layers and were trained on hundreds of thousands to millions of sequences. With only a few thousand ABA responsive genes available for training, using these DNN models from scratch is not feasible (over-parameterisation, high risk of overfitting and poor generalization). Furthermore, their complex architecture makes DNNs difficult to interpret and complicates the extraction of biologically relevant motifs across genes (39, 40). We therefore set-out to develop an interpretable CNN-based DL model to predict ABA responsiveness, starting from proximal promoter DNA sequences.

We mitigated the limited number of available genes for model training by implementing and comparing state-of-the-art augmentation techniques. The improved model performance when applying RC and EvoAug augmentation of proximal promoter sequences for training clearly reflects the general trend in DL that increased training data typically leads to better model performance. However, the contrast between classes is also important, as the performance for a model trained on all ABA UP genes is lower compared to a model trained on the 1200 most responsive genes (11). The 5′ UTR plays an important role in predicting ABA responsiveness, as models trained on genomic sequences including part of the 5’ UTR consistently outperform those that were trained on genomic sequences excluding them (9, 12, 14, 20). Shuffling the proximal promoter sequences eliminated model performance, underscoring that our model learned genuine motifs rather than simple nucleotide frequencies. Our model also outperformed a model trained to predict ABF binding, suggesting that it learned the cis-regulatory code beyond canonical ABF binding sites. Building upon the current state of the art of interpretable DL for genomic sequences (24, 25), we indeed showed that also other motifs, associated with different TF families (i.e., TALE, bHLH, NAC) are important for ABA-dependent gene expression regulation. For several of the identified TF families, we showed that homotypic motif clustering controls gene expression levels in response to ABA.

Previous studies already pointed towards the importance of distal interactions for gene expression regulation in complex plant genomes (41), but also in Arabidopsis (42). Using our model as a predictor we identified distal regulatory regions, beyond the proximal promoter used for model training, that were able to drive ABA-dependent transcriptional response *in planta*. The absence of ABF binding evidence in some of these distal regulatory regions corroborates our model learned ABA-specific regulatory code beyond canonical ABF binding sites. Distal regulatory regions were shown to be major drivers of context-specific gene regulation (41). Although the ABA-dependent gene expression was mainly induced in the elongation zone, particularly in the vasculature–consistent with previously reported ABA activity in the root (43)–we indeed observed notable variation in the cell-types in which these distal regulatory regions induced ABA responsiveness. One distal regulatory region, located in an intron, even induced ABA responsiveness in the leaves, and not in the root. These findings indicate that more subtle tissue and/or cell-type specific regulatory grammar was not captured by our model. Training cell-type-specific models on single-cell transcriptome data might provide a solution, but requires that sufficient informative genes are available. Furthermore, some genes were misclassified because the true regulatory sequences were not fully captured. Application of (sc)ATAC-Seq, plant self-transcribing active regulatory region sequencing (44) and/or promoter capture Hi-C (45) to define the input sequences for model training might therefore be important for future work as well (46).

## Materials and methods

### Processing expression data and extracting input region sequences

Supplementary tables reported DE were information downloaded from Sun and co-workers (17). The log2FC and adjusted p-values (q-values) of wild-type root untreated versus 3h ABA treatment were used to define our two classes. The genes were ranked using descending log2FC values and for a range of class sizes (n = 200, 400, 600, 800, 1000, 1200, 1400) the top n significantly (q-value≤0.05) DE genes were taken as the UP class. The NON class was subsequently defined as the equally-sized set of genes that had a q-value > 0.05 and log2FC closest to zero, without surpassing log2FC > 1 (*SI Appendix, Dataset S1*). For the dataset from Du and coworkers (21), the same class definitions were used.

Per gene, upstream input regions were defined relative to the TSS, and downstream (terminator) input regions relative to the TTS. Sequences 200, 500 and 1000 bp upstream of the TSS, with or without the first 100 bp downstream of the TSS (5’ UTR) were used as upstream input regions for model training. Sequences 100 and 500 bp downstream of the TTS, together with 100 bp upstream of the TTS (3’ UTR) were used as downstream regulatory regions for model training. Input regions were extracted from the Araport 11 GFF file downloaded from TAIR and these coordinates were converted to a bed file. “Bedtools getfasta” (bedtools v 2.30.0) (47) was used to extract the correct input sequences from the Arabidopsis Col-0 genome fasta file (Ensembl Plants release 59). Shuffled sequences were used as negative control.

In order to ensure that genes within the same family are not split between the training and test sets (12) genes were assigned to gene families using Orthofinder (version 2.2.3) (48). This information was used to split the genes in a train (80%), validation (10%), and test (10%) set, so that no gene family has genes in more than one of these sets.

### Data augmentation

Augmentation was implemented at train time to increase the number of training instances. For each sequence, the RC sequence was added and given the same label. Additionally, EvoAug (19) was applied with the default augmentation list, with max_aug_per_seq = 2 and hard_aug = True. Afterwards, to assess the effect of the augmentations included in the default augmentation list, a model was trained and tested for each separate augmentation. For the phylogenetic augmentation, we followed the framework of Duncan and coworkers (49). See *SI Appendix, Extended methods* for details on species sampling and orthology inference.

### Convolutional Neural Network architecture and training strategy

The CNN model consists of a convolutional layer with 64 kernels (size 12×4, stride 1), using one-hot encoded sequences as input, followed by a maximum pooling layer (window size 6, stride 6, dropout p = 0.25), a second convolutional layer with 64 kernels (size 6×1, stride 6, dropout p = 0.25) and a third convolutional layer with 32 kernels (size 6×1, stride 6, dropout p = 0.25). The output of the convolution is then flattened and passed to two dense layers of 256 and 128 units (dropout p = 0.25 in between dense layers), and finally to the prediction layer that has a single output unit. All convolutional and dense layers were followed by ReLU activation, except for the final prediction layer, which uses the sigmoid activation function (Fig. 1C). The model, with ∼90K parameters, was trained on both FW and RC sequences (see *SI Appendix, Extended methods*). To enforce RC invariance, post-hoc (i.e. after training) conjoining was used at inference; predictions of FW and RC sequences were averaged to obtain a gene’s prediction value (18). Genes with more than 50% probability were regarded as predicted UP and the rest were considered as predicted NON.

### ABA motif gold standard

A gold standard (GS) of motifs that are important for ABA response was determined to serve as a benchmark for the retrieval of important motifs by DeepExplainer and TF-MoDISco. Known TFBSs were collected from various databases and curated to remove redundant entries (see *SI Appendix, Extended methods*). A set of 2017 TFBSs were mapped on the proximal (−500,TSS,+100 bp) promoters of the genes of interest using Cluster-buster (CB), compiled on Sep 22, 2017 (50) and Find Individual Motif Occurrences (FIMO version 5.5.5) (51). Following (52), for both tools, default parameters were used, except for the CB gap parameter which was set to 20 and motif score to zero. The top motif matches of each motif were used to discard low-quality matches, retaining a maximum of 4000 motif matches for CB and a maximum of 7000 motif matches for FIMO. These mappings were subsequently used to perform TFBS enrichment analysis on the top1200 UP set (53). The list of significantly enriched TFBSs (q-value < 0.05) was filtered for TFs that are associated with an experimentally validated GO term related to ABA signaling, yielding 89 GS motifs. A motif was considered important if the corresponding TF was DE in response to ABA but lacked a GO annotation for ABA. We considered a TF DE when its log2FC was greater than 2 and its q-value < 0.05.

### DeepExplainer and TF-MoDISco

To perform motif analysis using DeepExplainer, 100 training instances were randomly chosen out of train, validation and test set to serve as the background for the DeepExplainer pipeline (24) (see *SI Appendix, Extended methods*). We used the same known TFBSs as used for defining the GS motifs to compare these TF-MoDISco motifs with known TFBS using TOMTOM (54), part of the TF-MoDISco pipeline (default settings).

### Experimental validation

Distal and intronic regions (best-scoring windows, see *SI Appendix, Extended methods*) beyond the proximal promoter, identified by the DL model as ABA cis-regulatory regions (prediction score > 0.5) were used to generate transgenic transcriptional reporter lines for *in planta* experimental validation. Selected best-scoring windows were fused in a vector to a minimal 35S promoter (m35S), the 5’-leader of tobacco mosaic virus (TMV), GUS, the HEAT SHOCK PROTEIN 18.2 (*HSP18*.*2*) terminator, and the NOPALINE SYNTHASE (*NOS*) terminator using Golden Gate cloning (see *SI Appendix, Table S7*). All constructs were verified by whole plasmid sequencing with Oxford Nanopore technology and were transformed into Arabidopsis plants using floral dipping (55) (see *SI Appendix, Extended methods*). Seedlings from independent transformations for each transgenic transcriptional reporter line were grown vertically for five days after germination at 22°C in continuous light conditions and then transferred to 1/2 MS plates containing 10 µM ABA (n = 5) for 17 hours or were left untreated (n≥3) and then harvested for GUS staining. The conditions for GUS staining were based on previously described protocols (56) (see *SI Appendix, Extended methods*). Seedlings were kept in X-Glc solution until staining was visible or up to 15 hours (overnight). GUS stained seedlings were mounted on slides using the imaging solution and imaged with a DIC microscope (Olympus BX51 DIC) at 100x magnification.

## Data, Materials, and Software Availability

Custom code used in this study is available from the GitHub repository (https://github.com/VIB-PSB/DeepDifE) and Zenodo (https://zenodo.org/records/15489116)

## Acknowledgments

This work was funded by the Ghent University Special Research Fund (BOF20/GOA/012) to M.M. and B.D.R., and the Research Foundation-Flanders (FWO) for ELIXIR Belgium (I000323N) to M.V.B. and S.T.

